# Male-benefit adaptation under sex-limited selection shaped by compensatory evolution in *Drosophila melanogaster*

**DOI:** 10.64898/2026.03.03.709222

**Authors:** Harshavardhan Thyagarajan, Mindy G. Baroody, Imran Sayyed, Joshua A. Kowal, Troy Day, Adam K. Chippindale

## Abstract

Intralocus sexual conflict (IaSC) results from opposing selection acting on traits with correlated expression between the sexes. We recently reported on a male-limited (ML) selection experiment in *Drosophila melanogaster* designed to investigate IaSC through a sex-limited evolution regime that theoretically resolves conflict in favour of males. However, this experiment did not universally or unambiguously improve male fitness, although female fitness declined as predicted. Here we examine sources of unintended selection: unusual genetic features of the breeding design and the special females used to enforce male-limited inheritance that may have complicated evolutionary outcomes. Specifically, we evaluate the effects of a foreign cytoplasm, genetically marked translocated autosomes, and a female-exposed Y chromosome derived from the clone-generator (CG) system, and the unique environment of sexual selection introduced by foreign CG females. We found that selected male fitness increased by 66% when expressed within the full ML genetic context, rising to over 100% when interacting with CG females. While there is no consistent fitness advantage in the “wild type” genetic background, there is a nearly significant trend of improved fitness with CG females (26% improvement). Further, outside this context, these males do not experience a fitness loss relative to controls, even showing a marginal gain of 6%. Uniformly, these gains were mediated by precopulatory traits: ML selection produced more attractive males with greater mating success and shorter mating latencies, while sperm competition remained unchanged. Intriguingly, ML-evolved males also exhibited reduced mate harm to females, contrary to the established narrative of escalating intersexual antagonism in this system. Dissecting individual components revealed significant fitness improvements associated with adaptation to the foreign cytotype (18%) and the female-exposed Y chromosome (33%), although responses varied across replicate populations. Moreover, when selected haplotypes were expressed together with the foreign cytotype in females, we observe a recovery in fitness. Together, these findings demonstrate extensive compensatory evolution to the ML selection environment, indicating that responses to the release of IaSC were shaped not only by sexually antagonistic selection but also by adaptation to the genetic manipulations and mating context inherent to the experimental design.

## INTRODUCTION

Gonochoric organisms experience fitness costs when sex-specific selection acts on shared traits, a phenomenon termed intralocus sexual conflict (IaSC). As sex-specific selection is common and traits are often genetically correlated between sexes, IaSC is expected to be widespread. It is predicted to influence: (1) maintenance of genetic variance for fitness through balancing selection, (2) evolution of sexual dimorphism, and (3) patterns of sex-biased gene expression and asymmetric selection on male- and female-biased genes.

IaSC has been studied using a variety of approaches: comparative anatomy, quantitative genetic analyses, genome-wide association, and experimental evolution with model organisms. *Drosophila melanogaster* features prominently, particularly with the analysis of intersexual genetic correlations (e.g., Chippindale *et al*. 2001), and “sex-limited” evolution (e.g., Rice 1998, Prasad *et al*. 2007) revealing evidence of a negative association between genetic variation favoured in one sex and effects upon the other. Sex-limited experimental evolution has served as a key experimental tactic within this body of work. These studies use cytogenetic systems that restrict chromosome transmission through a single sex (typically males), allowing haplotypes to respond exclusively to selection in that sex. If IaSC is present, fitness gains in the selected sex should come at the expense of the opposite sex through shifts in allele frequencies at sexually antagonistic loci.

The classic design for sex- (male-) limited selection in *Drosophila* is the “clone-generator” (CG) system reported by Rice (1996). In this breeding scheme, a combination of visible markers and genetic aberrations (a compound-X chromosome and translocated major autosomes) allow patrilineal transmission of the X-chromosome, matrilineal transmission of a Y chromosome, and easy sorting of paternal autosomal haplotypes that are transmitted from father to son (fig. S1a) whilst the maternally-derived vehicles of their transmission are discarded each generation. In other words, whole genomic haplotypes – “hemiclones” – can be cloned into many copies and fertilize eggs that develop into males. With large populations of genetically diverse hemiclones, selection can occur exclusively on male function, with the CG females used only for propagation.

Rice (1996) showed that ML-evolved haplotypes could rapidly adapt to their evolutionarily “arrested” CG mates, enhancing male reproductive success and competitive ability while also increasing harm to females, reflected in higher mortality—likely a collateral effect of post-copulatory sexual selection. Rice (1998) later demonstrated that these fitness gains were generalizable and not dependent on the use of CG females. Further, development time of females expressing ML haplotypes exhibited slower development, which was interpreted as an indication of IaSC. In a subsequent ML experiment, Prasad *et al*. (2007) would confirm the slower development of ML-expressing females but attribute the intralocus conflict more explicitly to the optimization of male development, as males too slowed down under ML-selection. Sexual dimorphism in development time and body size is highly conserved in this species and remained unchanged under ML-selection. Prasad *et al*. (2007) explained these changes as a shift in the direction of extant dimorphism, with both sexes displaying reduced growth rate (later adult eclosion and smaller body size), or “masculinization” of development. Later work (Bedhomme *et al*. 2008; Abbott *et al*. 2010) showed that these smaller, slower-developing males were more symmetrical, more male-like in phenotype, and more attractive to mates.

Important differences between these two experiments’ outcomes cropped up in their comparison, leaving key aspects of ML-selection responses unresolved. While Rice (1996, 1998) emphasized responses in post-copulatory traits such as sperm competition and mate harm, Prasad *et al*. (2007) and Jiang *et al*. (2011) did not corroborate these findings despite using the same LH_M_ base population and a larger experimental scale, suggesting possible changes in standing genetic variation with long-term domestication (cf. Collet *et al*. 2016). The widespread and predominant use of descendants of LH_M_ in the *Drosophila* sexual conflict literature suggested the need for investigation of other populations.

We recently reported on a new experiment using the ML-selection approach with different populations of *D. melanogaster* (Thyagarajan *et al*. 2025). In contrast with previous studies employing this technique, we found no consistent increase in male fitness of haplotypes subject to male-limited (ML) selection relative to their matched controls (MC). Males from ML lines did not uniformly outperform MC controls in mate choice trials, fecundity induction or sperm offense, although females expressing ML-evolved chromosomes displayed a substantial reduction in fitness, consistent with IaSC. Importantly, the experiments were conducted on animals removed from the selection treatment, which includes several genetic aberrations and mutant phenotypes. Our data therefore present mixed evidence for IaSC and motivate the present investigation into the impact of genetic artefacts and compensatory evolution in the ML-evolution protocol.

In evaluating potential genetic artefacts and compensatory responses in the ML protocol, we consider that CG females contribute more than a neutral transmission vehicle. Beyond passaging hemiclones, they create an evolutionarily arrested sexual-selection environment (*a la* Rice 1996) and impose a distinctive genomic and cytoplasmic background that can systematically shape selection outcomes. The translocated II–III autosome pair enforces co-segregation through aneuploid mortality and carries recessive visible markers. Earlier ML studies largely assumed that the absence of visible mutant phenotypes reflected general recessivity of autosomal alleles, without directly testing fitness effects (Rice 1996; Prasad *et al*. 2007). The compound-X (XXY) karyotype extends ML selection to the X chromosome by producing Y-bearing eggs. This reverses the usual parent-of-origin pattern of sex chromosomes in males, potentially inducing fitness effects by altering canalized paternal X-linked epigenetic effects. In addition, this introduces the direct effects of the CG Y chromosome, which experiences alternating male and female selection regimes. Despite limited coding content, the Y can exert broad regulatory effects (Chippindale & Rice 2001; Lemos *et al*. 2008; Jiang *et al*. 2010); and is known to have regulatory influences even when expressed in XXY females (Branco *et al*. 2017). Finally, CG females supply a foreign cytotype: a constraint that cannot be removed through backcrosses because the compound-X is inseparable from its cytoplasmic background. Strict maternal mitochondrial inheritance favours female-benefit variants, even when harmful towards male function (Cosmides & Tooby 1980, Frank & Hurst 1996, Gemmell *et al*. 2004, cf. Keaney *et al*. 2019). This is expected to result in local adaptation within populations through nuclear compensation for male function (Rand *et al*. 2004, Dowling *et al*. 2008, Ågren *et al*. 2019).

Here we report measures of competitive reproductive fitness (CRF), mating success, fecundity induction, sex-ratio drive, and sperm offense (P2). These traits were assayed in a fully factorial design across two genetic backgrounds: a complete ML “home-court” (HC) background retaining all selection features, and a control-derived “wild-type” (WT) background lacking them. Assays were conducted with both clone-generator (CG) females and control-derived competitors or females (Cr), thereby capturing the restricted sexual-selection environment of ML evolution. We also measure mate harm by selected and control males (in both genetic backgrounds) on CG females. Additionally, we study the fitness effects of the CG cytotype, Y chromosome and autosomes, each independent of the remaining artefacts in the genetic background.

## METHODS

### Source populations, Selection treatment

The source populations, selection protocols and controls are described in detail in Thyagarajan *et al*. (2025). Briefly, three replicate populations were derived from the control-old (CO) lines after 25 generations of adaptation to a 14-day life cycle. From each, a male-limited (ML) and a matched control (MC) population were established. Sex-limited selection on males was imposed through repeated crosses between ML males and clone-generator (CG) females (fig. S1a). CG females cause paternal transmission of the X chromosome to sons rather than daughters, supply a Y chromosome, and force co-segregation of the major autosomes (cII and cIII); we refer to this co-segregating X–II–III unit as a “target haplotype.” Competition among 1,050 target haplotypes for reproductive success with CG females each generation constituted one round of ML selection. Completely asexual propagation was avoided by allowing a small fraction (5%) of ML genomes to recombine as diploids in females (labelled recombination chamber “RC” females) (fig. S1b). The selection experiment ran for more than 80 generations.

### Experimental animals

We systematically investigated adaptation to the release of intralocus sexual conflict (IaSC), both with and without the artefactual features of the ML selection treatment, examining these features individually and in combination. The specific composition and production of test animals and competitors are described in full below. Briefly however, combined genetic effects were assessed in two “genetic backgrounds”: (i) a complete genetic “home court” (HC), retaining all of the ML selection genetic context, or (ii) a “wild-type” (WT) control-derived background lacking all these features. Individual component artefacts of the ML home court were tested in isolation to determine their independent contributions. The role of sexual co-evolution was examined using CG and control (Cr) females to evaluate the effects of co-evolved versus naive mating partners.

#### Home court (HC) males vs. Wild type (WT) males

All test males assayed carried target haplotypes [cI(X), II and III] from ML / MC populations. To place target (ML and MC) haplotypes under HC conditions, we crossed males from each treatment to CG females. Red-eyed sons from these crosses carry a haploid genome from the target genotype, combined with a CG derived (maternally inherited) Y chromosome, cytoplasm, and translocated autosomes (fig. S2a). The crosses conducted to produce WT males are described in detail in Thyagarajan *et al*. (2025). Briefly, we use MC derived males to sire test offspring with either MC females (producing WT MC test animals) or RC ML females. Male offspring from these crosses carry a haploid genome from the target genotype, combined with a control derived cytoplasm, (paternally inherited) Y chromosome, and translocated autosomes (fig. S2b).

#### C2lone generator (CG) and Control recessive (Cr) females

In addition, we take into consideration the arrested environment of sexual selection, by assaying test animals with CG females and control-derived females. For phenotypically marked control-derived competitors or females, we used lines backcrossed to have recessive eye colour markers (either “peach”, *p^p^*, or recessive brown *bw^1^*). Both competitor stocks are outbred and vigorous; here we refer to them collectively as “Cr”.

Across HC and WT backgrounds, males were evaluated with CG and Cr females for competitive reproductive fitness (CRF), mating success, mating latency, copulation duration, fecundity induction, sex-ratio drive, and sperm offense, while mate harm was assessed using CG females only.

#### Individual Artefacts - Males

To explore specific genetic artefacts, we combined target haploid genomes with individual CG-derived genetic elements while drawing the remaining genetic background from control animals. These test groups—termed “CG-cyto,” “CG-auto,” and “CG-Y”—carry a cytoplasm, autosome pair, or Y chromosome from the CG females, allowing us to isolate the effects of each artefact. For these assays, we measured only competitive reproductive fitness (CRF), in combination with Cr animals, focusing on the net consequences of each CG-derived genetic component.

The cytoplasm of the CG female, critically including elements like mitochondria, seemed to be inseparable from the CX (DX) chromosomes because both are maternally inherited. However, about 60 generations into the selection experiment, we fortuitously encountered a mutant female from a CG mother that had undergone a spontaneous breakdown of the conjoint X chromosome pair and was fertile. This rare event allowed us an opportunity to isolate the CG cytoplasm from the DX chromosome. Using this mutant female, we backcrossed the CG cytoplasmic background into a stock homozygous for translocated autosomes (carrying a *bwD* marker) and balancer X chromosomes (*FM7a).* Likewise, we also backcrossed the control MC cytotype into the same stock, creating two cyto-lines labelled FMTW-CG (carrying the CG cytotype) and FMTW-C (carrying the MC cytotype).

Female FMTW-CG flies were crossed to ML and MC males to capture haploid target genomes in female offspring. Daughters carrying these target haplotypes (balanced with FM and *bwD* translocated autosomes) were then crossed to MC males, to produce target male offspring carrying the CG cytoplasm, with control Y and autosomes (fig. S3a).

To produce CG-Y males, we first crossed MC males to CG (*bwD*) females. Sons from these crosses carry a CG derived Y chromosome, along with control derived autosomes. In parallel, we crossed ML and MC males to FMTW-C females, producing daughters carrying a target (ML/MC) haploid genome (balanced with FM and *bwD* translocated autosomes). These sons and daughters respectively are crossed to produce target male offspring carrying the CG Y chromosome, with control autosomes and cytoplasm (fig. S3b).

To produce CG-auto males, we first backcrossed the CG males to MC females to create a line homozygous for CG autosomes. We then used MC males to replace the CG derived Y chromosome while retaining the translocated autosomes. Males from this line were crossed to females homozygous for ML and MC target haplotypes to produce target male offspring carrying the CG autosomes, with control cytoplasm and Y chromosome (fig. S3c).

#### Individual Artefacts - Females

We express ML and MC haploid genomes in female flies along with a single artefact at time, to test for correlated responses to male counter-adaptations to the selection artefacts. These flies carried a CG derived cytoplasm with MC derived X and autosomes (CY-cyto), or CG translocated autosomes with an MC derived X and cytoplasm (CG-auto). The target females derived were tested for CRF.

To capture the CG cytotype, we backcrossed MC males to the FMTW-CG line, to replace the autosomes and one of the X chromosomes with MC derived chromosomes. In parallel, we crossed ML and MC males to CG (*bwD*) females. Sons from this cross are crossed to the backcrossed females to produce target female offspring carrying the CG cytoplasm, with control autosomes (fig. S4a).

To study CG autosomes, we backcrossed MC males to CG females to produce males homozygous for the translocated autosomes, in combination with a MC derived Y chromosome. These males were then crossed to females homozygous for ML and MC target haplotypes to produce target female offspring carrying the CG autosomes and a control cytoplasm (fig. S4b).

### Assays

The design used for CRF, mate choice and sperm competition assays are described in detail in Thyagarajan *et al*. (2025). They are described briefly below, with additional notes on any deviations from the original design made to accommodate the CG females.

#### Competitive Reproductive Fitness (CRF) assays

CRF was measured as competitive siring/damming success from mating arenas (ventilated ½ pint bottles) of 30 target animals combined with 30 Cr competitors of the same sex and 50 individuals of the opposite sex. Target animal output was measured by phenotyping the progeny using eye colour. In assays using CG females, we assess 10 such competition arenas for both MLs and MCs within each replicate; and collect all the eggs laid (typically yielding 1-2 vials of ∼80 viable individuals). For all other CRF assays, we assess 6 such competition arenas and collect 6 vials of ∼80 eggs.

#### Mate-choice assays

Female mate choice (or male-male competition) was assayed by combining a single female (CG or Cr) with a target male and Cr male. We identify the sire by phenotyping the progeny using eye colour. The resulting brood is also sexed and enumerated to test for fecundity initiation and sex ratio drive effects. Matings were observed to estimate mating latency and copulation duration. For each treatment (within each replicate pair of lines), we assess 50 such competition vials.

#### Sperm offense assays

Sperm offense (or P2) was assayed by serially mating CG or Cr females first to males that carried the same markers and subsequently to target males. P2 was measured as the proportion of target male progeny. For each treatment (within each replicate pair of lines), we assess 30 such competition vials.

#### Mate harm assay

Mate harm was assayed by measuring female mortality and productivity in response to being exposed to males. Following the crossing design, eggs of test animals are collected at densities of ∼80 larvae per vial. Test males werecollected on day 10-11. On day 12, 10 test males were combined in vials with 12 virgin CG females of the same age. Flies were held together for days 12-14, at which point the males were removed from the females using gentle anaesthesia. Female mortality was recorded daily for 7 days starting with the introduction of males, and the productivity of the females post mating was measured in the form of pupal counts.

### Statistical Analysis

Statistical analysis was conducted in R version 4.5.2 -- “Shortstop Beagle” (R Core Team 2025). Data was visually assessed for residual normality and using Bartlett’s test for heterogeneity. Where generalised linear models were used, residual dispersion was tested using the DHARMa package (Hartig 2022).

Linear mixed models were employed to analyse CRF. For HC and WT males, selection, genetic background, female and replicate population were used as fixed factors in the analysis to explain the proportion of offspring sired by the target individuals. To avoid pseudo-replication, contest was included as a random factor. Where significant interaction effects were identified, we employed planned contrasts between the levels of selection under each set of conditions with a corrected alpha value. For all the single artefact assays, selection and replicate were used as fixed factors, with contest used as a random factor. Where significant replicate differences were identified, we employed planned contrasts between the levels of selection in each replicate population pair with a corrected alpha value.

To study the mating success of target males in female mate choice trials, we used a generalized linear model with a binomial distribution. The proportion of mating successes for target animals was explained using selection, background, female, and replicate as fixed factors. Likewise, log(mating latency), mating duration, fecundity initiation (offspring count) and sex drive (offspring sex ratio) were analysed using a Gaussian distribution for tubes, with selection, background, female, and replicate as fixed factors. Where significant replicate differences were identified, we ran models with the same fixed factors within each replicate level with a corrected alpha value.

For sperm offense, the proportion of offspring sired by the second male was analysed in two steps, as if in a Hurdle model. First, to create a data transformation that enabled the analysis, we subtract the proportion of offspring sire from 1. This inverted measure is then separated in two parts – values of zero (where all the offspring are sired by the target male) and non-zero values. The number of zeroes attributable to each treatment was analysed as a generalized linear model with a binomial distribution and using selection, background, female, and replicate as fixed factors. The non-zero proportion is analysed using a Gaussian distribution of errors, with selection, background, female, and replicate as fixed factors.

For mate harm data, we first modelled the decline in survivorship of females after the males were removed (day3-day7) using a linear model with selection, background and replicate as fixed factors. This was based on a visual observation the rate of female mortality slowed down after the removal of the males. This model showed no effect of selection, background or replicate on female mortality. Based on this, we made the decision to model the female mortality using data from day 0-day 2 (period of male exposure). Again, we use a linear model with selection, background and replicate as fixed factors. Likewise, we use a linear model with selection, background and replicate as fixed factors to analyse productivity of the females mated to each treatment of males. Where significant interaction effects were identified, we employed planned contrasts between levels of selection under each set of conditions with a corrected alpha value.

## RESULTS

### HC and WT animals

#### Competitive reproductive fitness

Selection and genetic background both had strong effects on CRF (each p < 0.0001), with additional interactions between selection × background (p = 0.0364) and female type × background (p < 0.0001) (fig. 1b). Replicate had no effect and did not interact with other factors. The main model violated variance homogeneity (Bartlett’s p < 0.0001).

**Figure 1.**
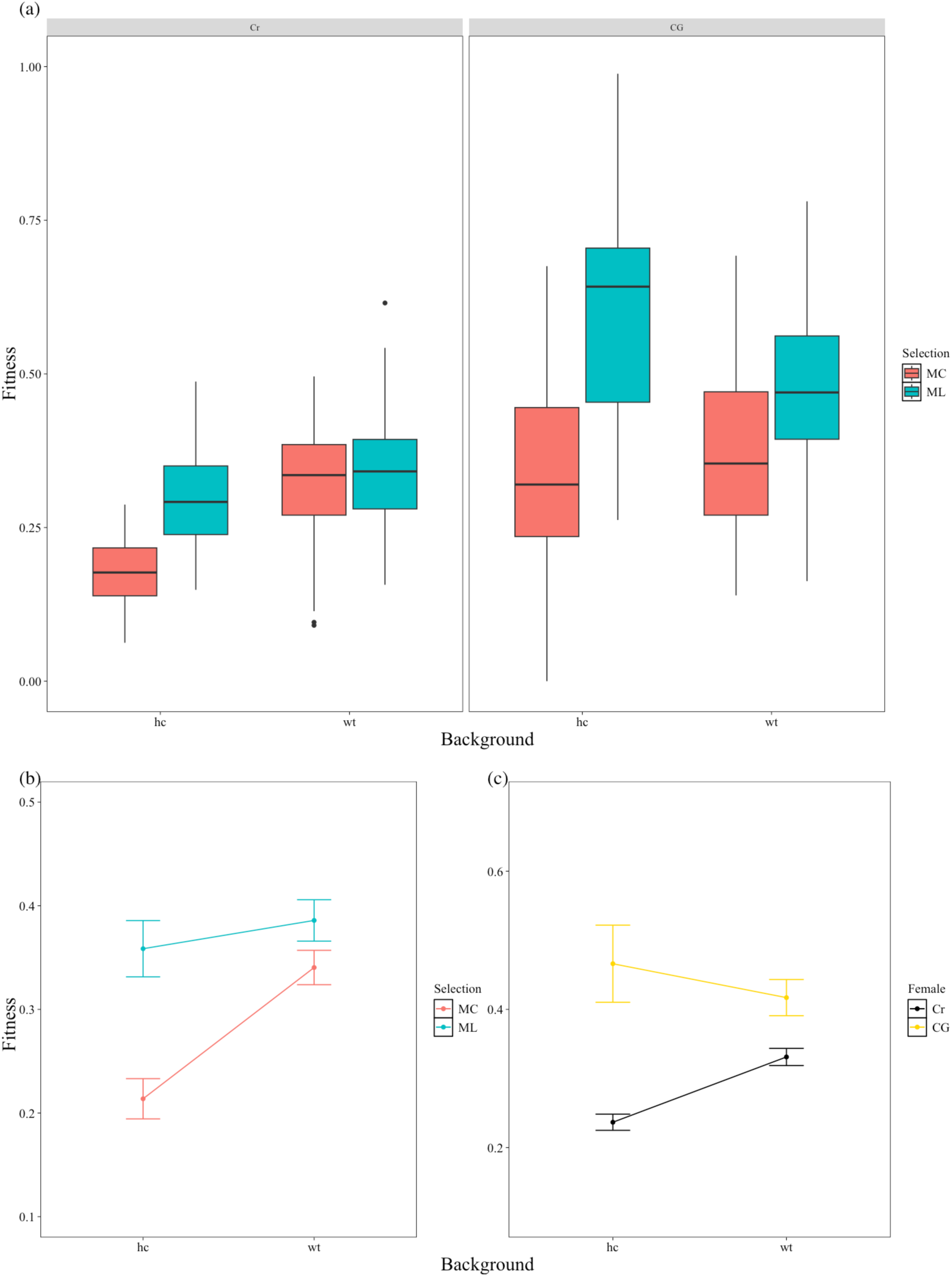
CRF of target males in HC (home court - ML like) & WT (wild type - MC like) genetic backgrounds, tested against (a) recessively marked control females and (b) clone generator females used in the ML selection experiment. Panels (c) & (d) highlight the interaction between (c) selection and genetic background and (d) female and genetic background.

As indicated by significant interactions, the fitness effects of the selection treatments were dependent on the context of female × background combination. Under home-court (HC) conditions with CG females, male-limited (ML) males had about twice the CRF of matched controls (MC) (0.224 ± 0.035 vs 0.110 ± 0.025, mean ± 1.96 SE). A similar pattern occurred in HC with Cr females, where ML males showed a 66% higher CRF (0.295 ± 0.015 vs 0.178 ± 0.010). Under wild-type (WT) conditions, ML males had about a 26% higher CRF with CG females (0.468 ± 0.038 vs 0.372 ± 0.033), although this effect did not meet the adjusted significance threshold (p = 0.0189 vs adjusted α = 0.0127). In contrast, differences were small with Cr females in WT conditions, where ML males showed only about a 6% relative increase (0.341 ± 0.018 vs 0.322 ± 0.017) (fig. 1a). Replicate effects were absent across all subsets.

#### Mate choice

We find significant effects of selection (p=0.0006), female (p=0.0144)), and replicate (p= 0.001) on the success of target males in mate choice trials. We find no effect of background or interaction effects. ML males outperformed MC males in mate choice trials, achieving on average about a 23% higher success rate across replicates (rep 1: 63.93% vs 51.58%; 3: 60.10% vs 48.62%; 5: 48.40% vs 40.98%) (fig. 2a).

**Figure 2.**
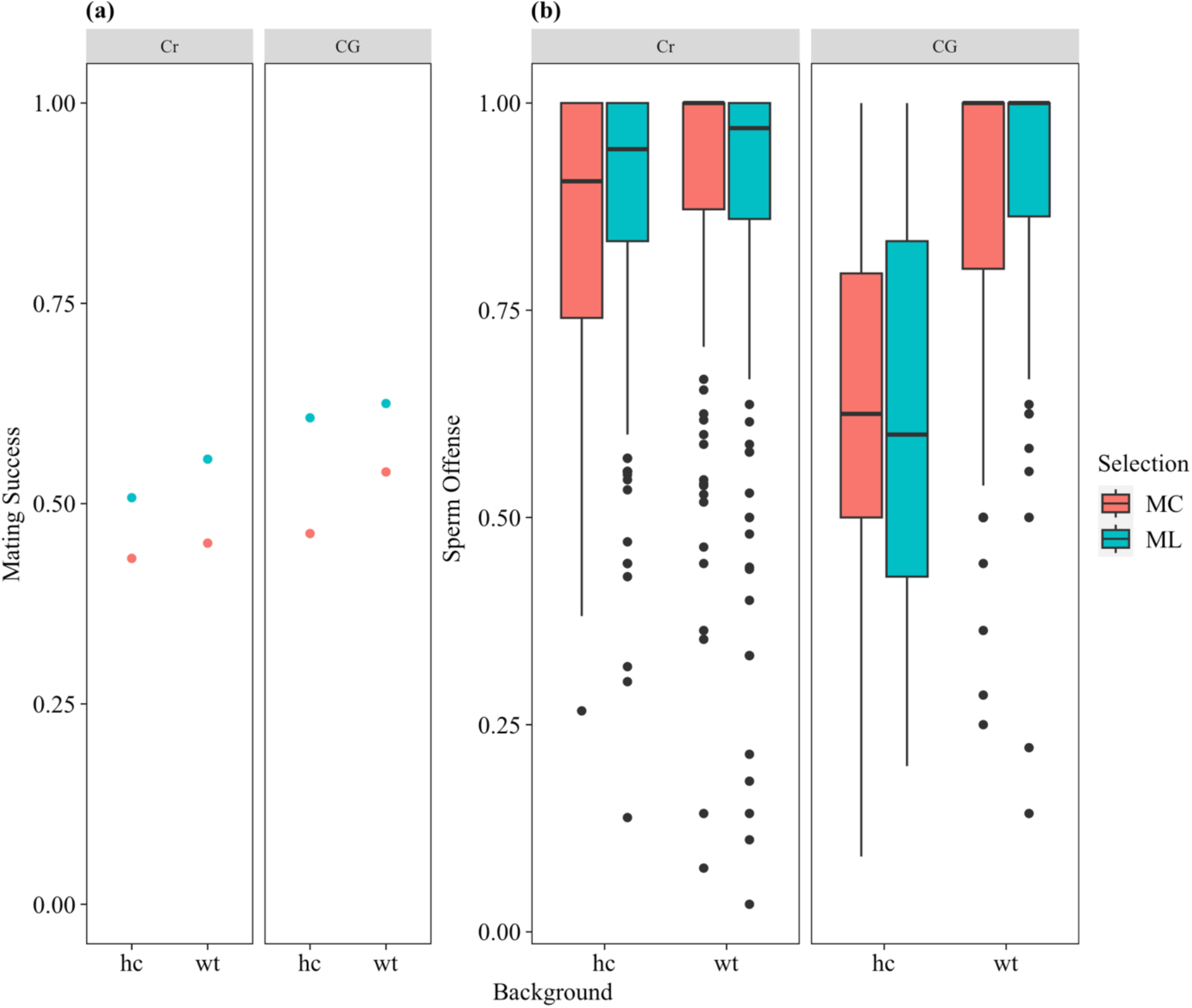
(a) The proportion of mating success of target males and (b) the proportion of offspring sired by target males as second mate (P2 - sperm offense). Target males from each genetic background (HC & WT) are tested against both kinds of females (recessive competitors - Cr, and clone generators - CG).

ML males also achieved matings faster, with 18% shorter latency than MC males across all conditions (14.05 ± 2.08 min vs 17.17 ± 2.89 min) (fig. 3a). Our analysis of mating latency reveals a significant effect of selection (p=0.0277) alone, with no effect of female, background, replicate or interactions on latency. Further, ML males mated longer than MC males did.

**Figure 3.**
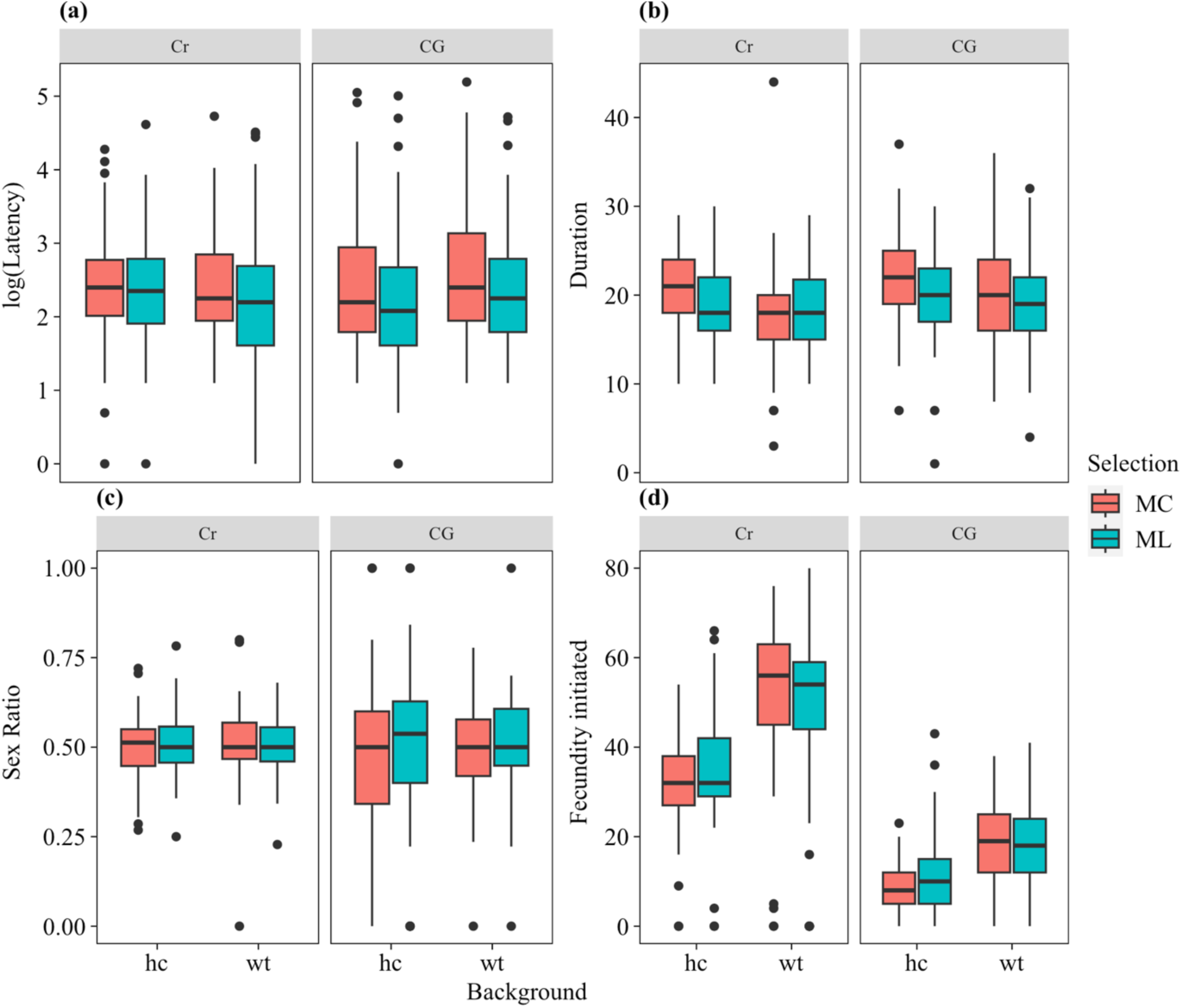
Measures of (a) mating latency, (b) mating duration, (c) brood sex ratios and (d) fecundity induction for target males from each genetic background (HC & WT), tested against both kinds of females (recessive competitors - Cr, and clone generators - CG).

Copulation duration was influenced by selection (p = 0.0167), background (p = 0.0007), female type (p = 0.0003), and replicate (p = 0.0197), with no interaction effects. Overall, ML males mated for about 4% shorter than MC males (19.76 ± 0.41 min vs 20.49 ± 0.43 min) (fig. 3b).

Across replicates, the proportional differences were similar: replicate 1 (20.07 ± 0.87 vs 20.57 ± 0.95 min), 3 (20.10 ± 0.68 vs 20.73 ± 0.83 min), and 5 (19.44 ± 0.73 vs 20.48 ± 0.73 min).

#### Fecundity induction, offspring sex ratio

Fecundity induction showed no effect of selection (27.80 ± 2.20 vs 27.18 ± 2.46) (fig. 3d), although background (p < 0.0001), female (p < 0.0001), replicate (p = 0.013), and the female × background interaction (p < 0.0001) were significant. Offspring sex ratio also showed no effect of selection, with ML males producing broods with a sex ratio of 0.505 ± 0.011 compared to 0.494 ± 0.011 for MC males (fig. 3c).

#### Sperm offense

Across both steps of our sequential model to analyse sperm offense, we find no effect of selection or significant interaction with selection on P2. In both steps, we find significant effects of background, female and an interaction between female and background. Overall, MLs display a P2 of 0.848 +/- 0.0169 compared to and MC 0.825 +/- 0.0175 (fig. 2b).

#### Mate harm

Female mortality during male exposure was 25–45% lower for ML males than MC males, depending on background (HC: 2.40 ± 0.37 vs 3.20 ± 0.53, ∼25% lower; WT: 1.35 ± 0.33 vs 2.37 ± 0.47, ∼43% lower; selection p < 0.0001) (fig. 4a). Replicate and interaction effects were not significant.

**Figure 4.**
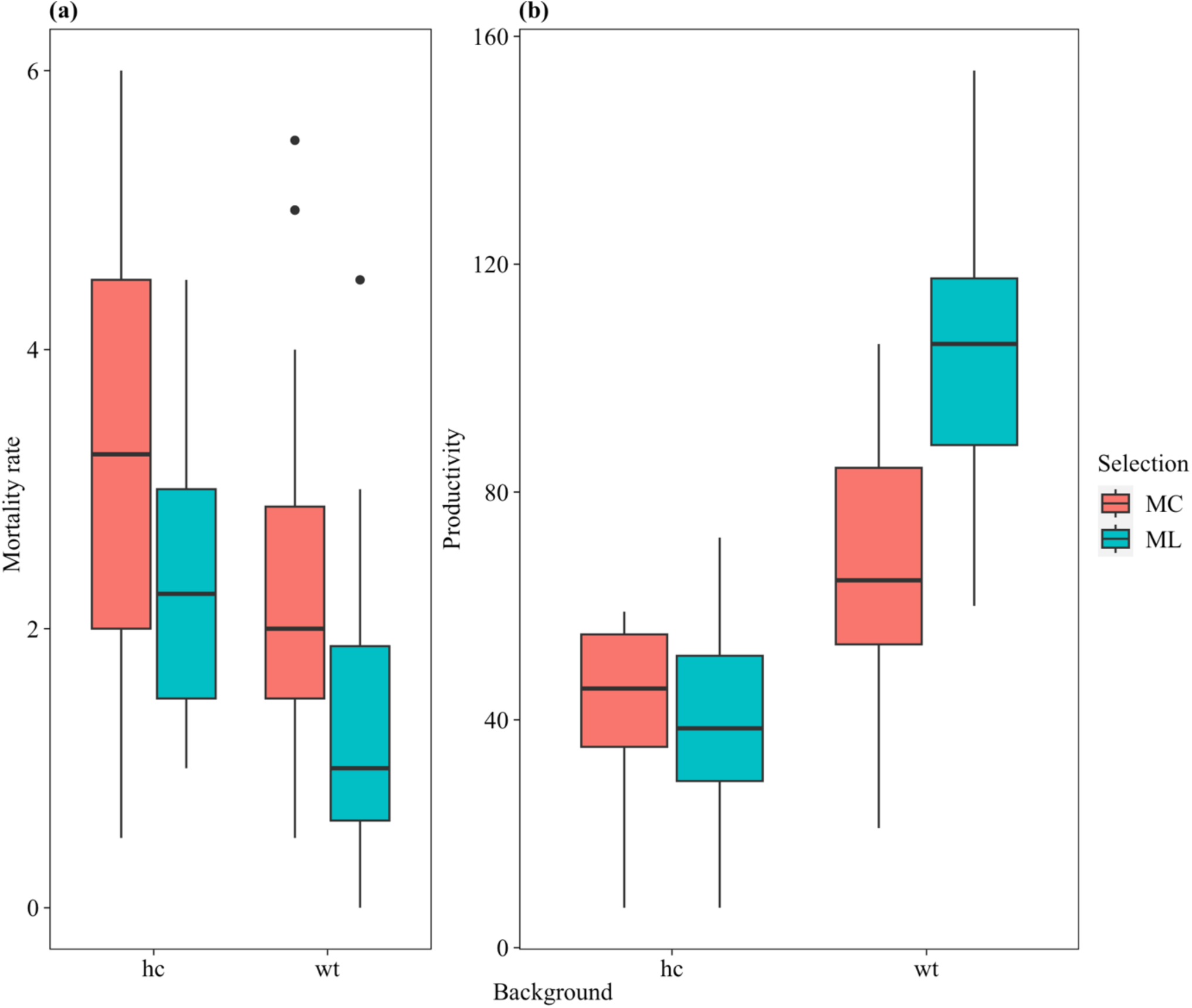
Clone generator (CG) female (a) mortality per diem during male exposure and (b) pupal productivity upon exposure to target males from both home court (HC) and wild type (WT) genetic backgrounds.

Productivity differences depended on background. In the HC background, we found no significant effect of selection (40.0 ± 5.36 vs 43.5 ± 4.76). In the WT background, ML males produced about 53% higher productivity than MC males (104.0 ± 7.54 vs 68.0 ± 8.33, selection p < 0.0001) (fig. 4b). Replicate and interaction effects were not significant in either background.

### Single artefact animals – CRF

#### CG cytoplasm

For males with the CG cytotype, selection significantly increased CRF by about 18% (ML: 0.360 ± 0.019 vs MC: 0.304 ± 0.016, p = 0.0225) (fig. 5b), with no effect of replicate or interaction. For females with the CG cytotype, CRF was influenced by replicate (p < 0.0001) and the selection × replicate interaction (p = 0.0334), while selection overall was marginal (p = 0.0517). Examined by replicate, selection had a significant effect only in replicate 5 (∼9% lower CRF in ML females: 0.516 ± 0.014 vs 0.569 ± 0.015, p = 0.0004), but not in replicates 1 (0.488 ± 0.017 vs 0.488 ± 0.016) or 3 (0.494 ± 0.017 vs 0.497 ± 0.014) (fig. 5d).

**Figure 5.**
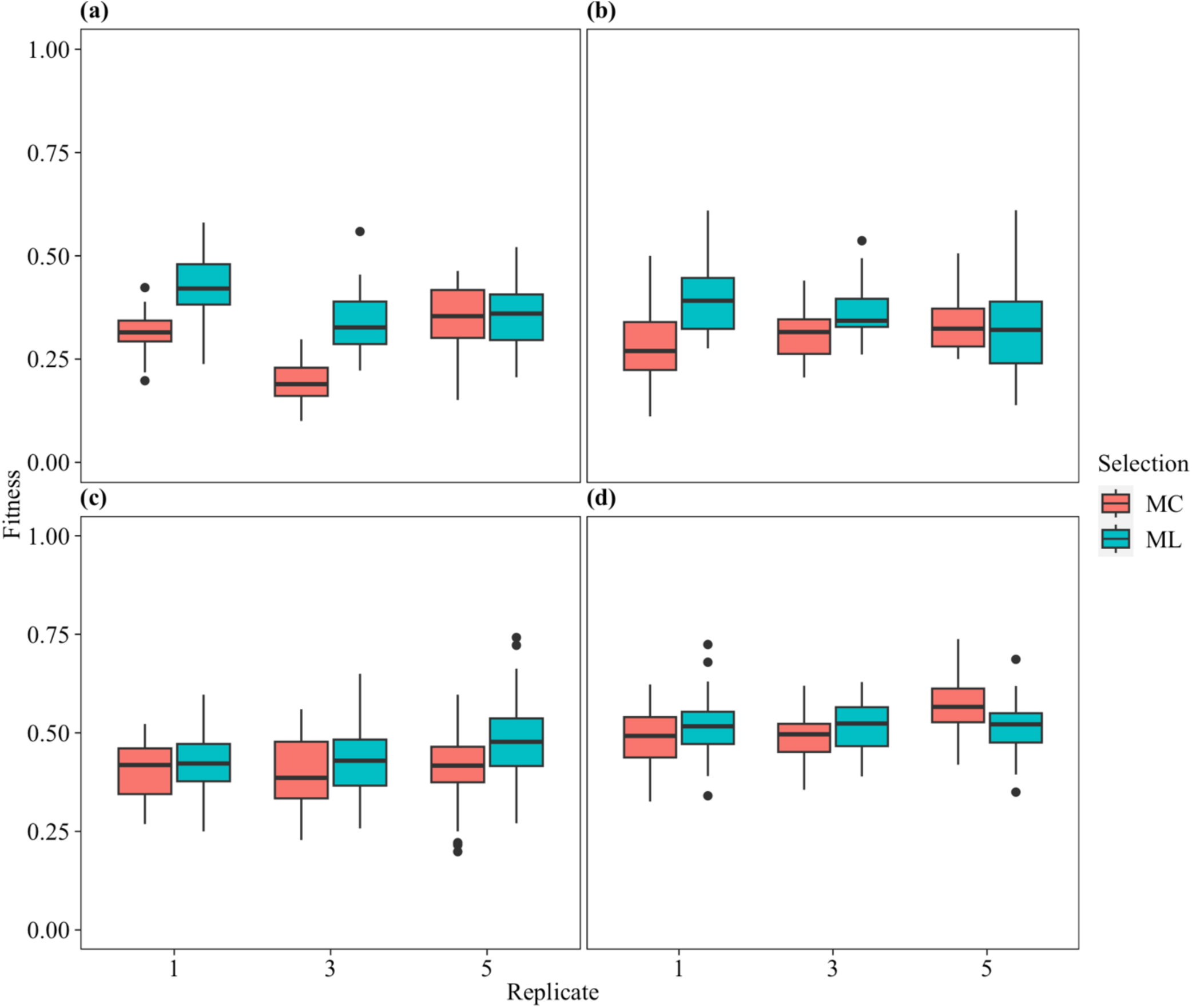
Competitive reproductive fitness (CRF) of target males with a completely control genetic background except for (a) CG-Y chromosome, (b) CG-cytoplasm, (c) CG-autosomes; and CRF of target females with a completely control genetic background except for (d) CG-cytoplasm.

#### CG Y

For males with the CG Y chromosome, CRF was significantly affected by selection (p < 0.0001), replicate (p < 0.0001), and their interaction (p = 0.0071). Overall, ML males displayed about a 33% higher CRF than MC males (average across replicates: 0.376 vs 0.284). When examined by replicate, selection significantly increased CRF in replicates 1 (0.431 ± 0.029 vs 0.312 ± 0.017, p = 0.0011) and 3 (0.340 ± 0.026 vs 0.198 ± 0.020, p = 0.0008), but not in replicate 5 (0.357 ± 0.029 vs 0.342 ± 0.033, p = 0.7065) (fig. 5a).

#### CG Autosomes

We find no effect of selection, replicate, or interaction between selection and replicate on CRF of males with target haplotypes paired with CG autosomes. ML males displayed an overall CRF of 0.447 +/- 0.019, compared to MC males at 0.405 +/- 0.016 (fig. 5c).

For females with CG autosomes, CRF was influenced by selection (p < 0.0001), replicate (p = 0.013) and the selection × replicate interaction (p < 0.0001). Overall, ML females displayed about a 22% lower CRF than MC females (average across replicates: 0.169 vs 0.215). When examined by replicate, selection significantly decreased CRF in replicates 3 (0.196 ± 0.017 vs 0.151 ± 0.017, p = 0.0005) and 5 (0.247 ± 0.025 vs 0.154 ± 0.014, p = 0.0004), but not in replicate 1 (0.201 ± 0.020 vs 0.201 ± 0.019, p = 0.96).

## DISCUSSION

Most of the selection experiments on sexual conflict in *Drosophila* using the male-limited (ML) selection protocol developed by Rice (1996, 1998), or variants, were conducted on a single base population of fruit flies, LH_M_, leaving us wondering about the generality of the earlier results. We recently reported (Thyagarajan *et al*. 2025) that a new application of the ML selection protocol provided mixed evidence for the resolution of intralocus sexual conflict in three newly investigated populations of *D. melanogaster*. We found a clear signal of reduced fitness in females expressing male-limited (ML) evolved chromosomes, and males that exhibited increased fitness in some assays but not others, potentially reflecting adaptation to unintended sources of selection – unnatural genetics, such as paternal X and maternal Y sex chromosomal inheritance, novel cytoplasmic properties (notably the mitochondria carried by the special DX females), and the translocated autosomes (t2;3). Here we dissect the evolutionary response to the release of intralocus sexual conflict (IaSC) across conditions that retain or remove the artefactual features of the ML selection protocol, examined both individually and in combination. We show that these populations exhibit a substantial response to selection, whose expression depends strongly on both male genetic background and the female context in which males are evaluated.

The magnitude of adaptive response was considerable: we found that ML males were approximately twice as fit as controls (+103.6%) when tested with all of the features of their selection protocol – in their genetic “home court”, competing for mating with the special “clone-generator” (CG) females they had potentially become specialized upon. With control females that these males could not be specially adapted to, ML-evolved males in their “home court” were about 66% fitter than control males were. While we do not find fitness advantage for ML selected males in the “wild type” genetic background, there is a nearly significant trend of improved fitness with CG females (26% improvement). Therefore, in addition to imposing costs on females, ML selection generates fitness gains in the selected sex. These fitness gains are largely context-specific, reflecting adaptation to the artefactual costs of the breeding design and clone-generator mating environment. Further, outside this context, these males do not experience a fitness loss relative to controls and even display a marginal gain of 6%.

We first tried to partition the fitness improvement seen in ML-selected males into the sphere of sexual selection it occurred in. The fitness increase in selected males was primarily explained by increased competitive mating success, with no significant changes between selected and control lines in fecundity induced in mates, brood sex ratio or sperm offense (P2). The evolved response was thus generally found to be in the realm of pre-copulatory sexual selection rather than in adaptation to the female post-mating reproductive response. This result aligns with earlier work on LH_M_-derived flies (Bedhomme *et al*. 2008; Jiang *et al*. 2011) which reported increased mating success of ML-selected males, and no response in post-copulatory sexual selection. However, these results are in marked contrast to Rice (1998) whose work implicated a response in the realm of sperm competition and ejaculate-female interactions. The overall interpretation of the earlier experimental work from the Chippindale group was one of enhanced male attractiveness, perhaps involving phenotypic masculinization and increased symmetry of ML-evolved males (Bedhomme *et al* 2008, Abbott *et al*. 2010). By contrast, it is unintuitive for a sex-limited trait like sperm competition to respond further to male-limited selection except via correlated effects of improved male condition, which might explain why post-copulatory traits showed no change here or in Jiang *et al*. (2011), contrary to Rice (1998).

Rice (1996) reported increased mate harm by ML-evolved males, detectable in mortality increases from even a single mating and thus suggestive of an ejaculate-female interaction rather than sustained harassment. Here we find the opposite: when mating with the CG females, ML-selected males caused substantially less mate harm than controls in both genetic backgrounds (HC and WT). During the selection process, CG females were so frail that we often had difficulty maintaining the ML populations in the face of their rapid mortality. Since we did not detect the kinds of fertility-inducing effects or advantages in sperm competition that have been suggested to drive male ejaculate toxicity in this species, our results raise an intriguing question about the nature of intersexual interactions: can selection favour reduced mate harm in the absence of monogamy or kin selection? It appears that less harming males can be favoured by selection, at least when females are feeble - as in the conditions we maintained our lines.

The ML design makes use of reversed sex-chromosome transmission, with paternal inheritance of the X and maternal inheritance of the Y. Like previous users of the CG system, we originally assumed that the Y chromosome was inert and could not be modified by maternal transmission. However, our finding of strong compensatory evolution for the impacts of the CG system forced us to consider the possibility that the Y chromosome could be modified by carriage in females through selection or possibly modification during transmission through the egg. We found evidence that the presence of this “feminized” Y chromosome results in a mating advantage for selected males, relative to MC control males also carrying it, even in the WT background and in competition for mating with the Cr marker stock females (where it was not expected based on competitive reproductive fitness data).

In addition to adaptation to feminized Y chromosomes, our results indicate that the cytoplasmic background of clone-generator (CG) females also influences male performance. ML haplotypes experienced this cytotype throughout selection, whereas MC haplotypes encountered it only when expressed experimentally, creating a natural contrast in prior exposure. Among potential cytoplasmic factors, we excluded infection by *Wolbachia* using PCR assays (data not shown), and we observed no evidence of the female-biased sex-ratio distortion typical of parasitic endosymbionts such as *Spiroplasma*. With endosymbiont effects excluded, the mitochondrial (mt) genome remains the most plausible source of the cytoplasmic effect. This particular cytotype can be traced matrilineally to the origin of the compound-X chromosome in the 1940s (Muller 1943), raising the possibility that it is substantially diverged from the mitochondria present in the outbred Ives-derived populations used in our laboratory. Consistent with this speculation, recent sequencing shows that CG females carry a mitochondrial genome resembling the OreR/w¹¹¹⁸ lineage and distinct from that of the Ives-derived stocks used herein (F. Camus, pers. comm.; experiments in progress). Reductions in male fitness under nuclear–mt mismatch are widely interpreted as evidence for the “mother’s curse.” In our system, such effects would most plausibly arise in MC haplotypes exposed to the clone-generator cytotype under home-court conditions, where their nuclear genomes encounter a foreign mitochondrial background for the first time. This predicts a transient decline in MC male performance relative to both wild-type benchmarks and ML haplotypes that have undergone compensatory evolution in this context. Such a mismatch may underlie the early-generation (“CDX”) fitness decline reported in male-limited X-chromosome work (Abbott *et al*. 2013), although the mitochondrial genotype of the LHm base population relative to OreR/w¹¹¹⁸ remains unknown. We therefore suggest a double mother’s curse – mitochondria and feminized Y chromosomes – resulting from the CG females: the mc hurts the MCs.

Previously, we showed that female fitness of selected haplotypes declined under WT conditions (Thyagarajan *et al* 2025). Here we ask if compensatory adaptation in the males takes advantage of sex-limited selection to improve males through routes that are indifferent (sex-limited), beneficial, or antagonistic to female fitness. When the cytotype was isolated and tested, we found a recovery of fitness in females expressing the selected haplotypes relative to control females, suggesting that the nuclear response to the foreign cytotype produces sexually concordant benefits. In this sense, our results support nuclear-mt coadaptation more broadly, rather than the mother’s curse, where defective mt specifically afflict male function. By contrast, we detected no comparable compensatory response to the translocated autosomes: these elements did not significantly improve male performance, nor did they alter female fitness, with ML females remaining less fit than MC females, as predicted under classical IaSC expectations.

Compensatory effects were not entirely consistent across replicate populations, with variation in which mechanisms allowed fitness to improve in the ML genetic home court. For example, while selected males in population 5 show little improvement over controls with both the CG cytoplasm and CG Y chromosome, we find clear improvements in both cases for populations 1 and 3. Likewise, while females with selected haplotypes carrying CG cytotype show fitness recovery in replicates 1 and 3, such a recovery is not observed in replicate 5. The strongest improvement for replicate 5 is seen in the performance with the translocated (t2;3) autosomes of the CG system. While we find no significant effect of selection, overall, when assessing the effect of the CG autosomes on fitness, we observe that the selected animals in replicate 5 show a trend of improved fitness relative to their paired controls under these conditions. Taken together, the 3 replicate populations appear to follow different routes in compensatory evolution The compensatory adaptation displayed in the selection experiment raises concerns about previous experimental work conducted using the CG system for sex-limited evolution. CG flies have been used as dams in breeding designs to conduct hemiclonal analyses (Chippindale *et al*. 2001; Gibson *et al*. 2002; Byrne & Rice 2005; Lew & Rice 2005; Lew *et al*. 2005; Linder & Rice 2005; Pischedda & Chippindale 2006; Long & Rice 2007; McKean *et al*. 2008; Rode & Morrow 2009; Innocenti & Morrow 2010; Mallet *et al*. 2011; Tennant *et al*. 2014; Hill *et al*. 2017; Ruzicka *et al*. 2019; Abbott *et al*. 2020) and male-limited selection experiments (Rice 1996; Prasad *et al*. 2007). It is important to note that most of these experiments (all but McKean *et al*. 2008, Mallet *et al*. 2011 & Tennant *et al*. 2014) work with target haplotypes derived from the LH_M_ stock population, which may respond differently to artefactual selection pressures described herein. In different iterations, authors have taken various level of precautions against possible artefacts, ranging from Rice (1998) where everything but the CG Y chromosome was controlled for, to Rice (1996) where none of these artefacts were controlled for. A common theme running through all these studies is the use of test animals with CG derived cytotypes and Y chromosomes, frequently inherited directly from the dam. In this work we find that both can introduce fitness effects, differentially affecting selected and control animals in selection experiments, potentially distorting trait variance in hemiclonal analyses.

## Author contributions

HT, AKC and TD developed the overall theme of the study; HT and AKC designed the experiments; HT, AKC, IS, MGB, & JAK carried out the experiments; HT analysed the data; HT wrote the first draft of the paper; HT & AKC contributed to the final draft and revisions.

## Conflict of interest statement

The authors declare no competing interests.

## Acknowledgments

Several undergraduate students were involved in lab maintenance and data collection during the period of this study. We thank Avery Want, Halle McNamara, Ronni Prince, Katelyn Viau, Alex Montenegro-Monreal, and Simmal Grewal for their efforts. Funding for the work was from NSERC Discovery grants to AKC & TD.

## Data Accessibility Statement

## Notes

### Competing Interest Statement

The authors have declared no competing interest.

## REFERENCES

Abbott, J. K., Bedhomme, S., & Chippindale, A. K. (2010). Sexual conflict in wing size and shape in Drosophila melanogaster. Journal of Evolutionary Biology, 23(9), 1989–1997.

Abbott, J.K., Innocenti, P., Chippindale, A.K. & Morrow, E.H. (2013). Epigenetics and Sex-Specific Fitness: An Experimental Test Using Male-Limited Evolution in Drosophila melanogaster. PLOS ONE, 8, e70493.

Abbott, J.K., Chippindale, A.K., Morrow, E.H. (2020). The microevolutionary response to male-limited X-chromosome evolution in *Drosophila melanogaster* reflects macroevolutionary patterns. J Evol Biol; 33: 738–750.

Ågren, J. A., Munasinghe, M., & Clark, A. G. (2019). Sexual conflict through mother’s curse and father’s curse. Theoretical population biology, 129, 9–17.

Bedhomme, S., Prasad, N.G., Jiang, P.P. & Chippindale, A.K. (2008). Reproductive Behaviour Evolves Rapidly When Intralocus Sexual Conflict Is Removed. PLOS ONE, 3, e2187.

Branco, A. T., Brito, R. M., & Lemos, B. (2017). Sex-specific adaptation and genomic responses to Y chromosome presence in female reproductive and neural tissues. *Proceedings*. Biological sciences, 284(1869), 20172062.

Byrne, P. G., & Rice, W. R. (2005). Remating in Drosophila melanogaster: an examination of the trading-up and intrinsic male-quality hypotheses. Journal of evolutionary biology, 18(5), 1324–1331.

Chippindale, A.K., Gibson, J.R. & Rice, W.R. (2001). Negative genetic correlation for adult fitness between sexes reveals ontogenetic conflict in *Drosophila*. Proceedings of the National Academy of Sciences, 98, 1671.

Collet, J. M., Fuentes, S., Hesketh, J., Hill, M. S., Innocenti, P., Morrow, E. H., … & Reuter, M. (2016). Rapid evolution of the intersexual genetic correlation for fitness in *Drosophila melanogaster*. Evolution, 70(4), 781–795.

Connallon, T. & Matthews, G. (2019). Cross-sex genetic correlations for fitness and fitness components: Connecting theoretical predictions to empirical patterns. Evolution Letters, 3, 254–262.

Cosmides, L. M., & Tooby, J. (1981). Cytoplasmic inheritance and intragenomic conflict. Journal of theoretical biology, 89(1), 83–129.

Dowling, D.K., Friberg, U., Lindell, J., 2008. Evolutionary implications of non-neutral mitochondrial genetic variation. Trends Ecol. Evol. 23, 546–554.

Frank, S., & Hurst, L. (1996). Mitochondria and male disease. Nature 383, 224

Gemmell, N. J., Metcalf, V. J., & Allendorf, F. W. (2004). Mother’s curse: the effect of mtDNA on individual fitness and population viability. Trends in ecology & evolution, 19(5), 238–244.

Gibson, J.R., Chippindale, A.K. & Rice, W.R. (2002). The X chromosome is a hot spot for sexually antagonistic fitness variation. Proceedings of the Royal Society of London. Series B: Biological Sciences, 269, 499–505.

Hartig, F. 2022. “DHARMa: Residual Diagnostics for Hierarchical (Multi-Level / Mixed) Regression Models.” R package version 0.4.6

Hill, M.S., Ruzicka, F., Fuentes, S., Collet, J.M., Morrow, E.H., Fowler, K., et al. (2017). Sexual antagonism exerts evolutionarily persistent genomic constraints on sexual differentiation in *Drosophila melanogaster*. bioRxiv, 117176.

Frank, S. A., & Hurst, L. D. (1996). Mitochondria and male disease. Nature, 383(6597), 224.

Innocenti, P. & Morrow, E.H. (2010). The Sexually Antagonistic Genes of Drosophila melanogaster. PLOS Biology, 8, e1000335.

Jiang, P. P., Hartl, D. L., & Lemos, B. (2010). Y not a dead end: epistatic interactions between Y-linked regulatory polymorphisms and genetic background affect global gene expression in Drosophila melanogaster. Genetics, 186(1), 109–118.

Jiang, P. P., Bedhomme, S., Prasad, N. G., & Chippindale, A.K. (2011). Sperm competition and mate harm unresponsive to male-limited selection in Drosophila: an evolving genetic architecture under domestication. Evolution; international journal of organic evolution, 65(9), 2448–2460.

Keaney, T. A., Wong, H. W. S., Dowling, D. K., Jones, T. M., & Holman, L. (2020). Mother’s curse and indirect genetic effects: Do males matter to mitochondrial genome evolution? Journal of evolutionary biology, 33(2), 189–201.

Kidwell, J.F., Clegg, M.T., Stewart, F.M. & Prout, T. (1977). Regions of stable equilibria for models of differential selection in the two sexes under random mating. Genetics, 85, 171–183.

Lemos, B., Araripe, L. O., & Hartl, D. L. (2008). Polymorphic Y chromosomes harbor cryptic variation with manifold functional consequences. *Science (New York*, N.Y.), 319(5859), 91–93.

Lew, T.A. & Rice, W.R. (2005) Natural selection favours harmful male *Drosophila melanogaster* that reduce the survival of females Evol Ecol Res 7: 633–641

Linder, J. E. & Rice, W. R. (2005). Natural selection and genetic variation for female resistance to harm from males. Journal of evolutionary biology, 18(3), 568–575.

Long, T.A.F. & Rice, W.R. (2007). Adult locomotory activity mediates intralocus sexual conflict in a laboratory-adapted population of Drosophila melanogaster. Proceedings of the Royal Society B: Biological Sciences, 274, 3105–3112.

Mallet, M.A. & Chippindale, A.K. (2011). Inbreeding reveals stronger net selection on Drosophila melanogaster males: implications for mutation load and the fitness of sexual females. Heredity, 106, 994–1002.

McKean, K.A., Yourth, C.P., Lazzaro, B.P. et al. (2008). The evolutionary costs of immunological maintenance and deployment. BMC Evol Biol 8:76.

Muller, H. J. (1943). A stable double X chromosome. Drosophila Information Service, 17, 61–62.

Pischedda, A. & Chippindale, A.K. (2006). Intralocus Sexual Conflict Diminishes the Benefits of Sexual Selection. PLOS Biology, 4, e356.

Prasad, N.G., Bedhomme, S., Day, T. & Chippindale, A.K. (2007). An Evolutionary Cost of Separate Genders Revealed by Male-Limited Evolution. The American Naturalist, 169, 29–37.

R Core Team. (2025). R: A language and environment for statistical computing (Version 4.5.2)

Rand, D. M., Haney, R. A., & Fry, A. J. (2004). Cytonuclear coevolution: the genomics of cooperation. Trends in ecology & evolution, 19(12), 645–653.

Rice, W.R. (1996). Sexually antagonistic male adaptation triggered by experimental arrest of female evolution. Nature, 381, 232–234.

Rice, W.R. (1998). Male fitness increases when females are eliminated from gene pool: Implications for the Y chromosome. Proc. Natl. Acad. Sci., 95, 6217–6221.

Rode, N.O., and Morrow, E.H. (2009), An examination of genetic variation and selection on condition in *Drosophila melanogaster* males. Entomologia Experimentalis et Applicata, 131, 167–177.

Ruzicka, F., Hill, M.S., Pennell, T.M., Flis, I., Ingleby, F.C., Mott, R. et al. (2019). Genome-wide sexually antagonistic variants reveal long-standing constraints on sexual dimorphism in fruit flies. PLOS Biology, 17, e3000244.

Thyagarajan, H., Sayyed, I., Baroody, M. G., Kowal, J. A., Day, T., & Chippindale, A. K. (2025). Mixed evidence for intralocus sexual conflict from male-limited selection in *Drosophila melanogaster*. *Journal of Heredity*, esaf072.

