## Supplementary material for "Male-benefit adaptation under sex-limited selection shaped by compensatory evolution in *Drosophila melanogaster*": S2a

Intralocus sexual conflict obscured through compensatory adaptation to artefacts in a male-limited selection experiment in *Drosophila melanogaster*.

**Table S1.** The results of the fully factorial ANOVA fit for the male CRF.

| *Response* | *Variable* | *df* | *F* | *P* |
|---|---|---|---|---|
| Fitness | Selection | ***1*** | ***39.0051*** | ***<0.0001*** |
| | Replicate | ***2*** | ***0.4380*** | ***0.65*** |
| | Female | ***1*** | ***0.7970*** | ***0.37*** |
| | Bg | ***1*** | ***182.9934*** | ***<0.0001*** |
| | Sel:Rep | ***2*** | ***0.2625*** | ***0.77*** |
| | Sel:Fem | ***1*** | ***1.4094*** | ***0.24*** |
| | Sel:Bg | ***1*** | ***4.3954*** | ***0.0376*** |
| | Fem:Bg | ***1*** | ***32.1444*** | ***<0.0001*** |
| | Fem:Rep | ***2*** | ***1.9557*** | ***0.14*** |
| | Bg:Rep | ***2*** | ***0.0473*** | ***0.95*** |
| | Sel:Fem:Bg | ***1*** | ***1.4517*** | ***0.23*** |
| | Sel:Bg:Rep | ***2*** | ***0.9330*** | ***0.39*** |
| | Sel:Fem:Rep | ***2*** | ***0.0503*** | ***0.95*** |
| | Fem:Bg:Rep | ***2*** | ***0.1795*** | ***0.84*** |
| | Sel:Fem:Bg:Rep | ***2*** | ***2.4533*** | ***0.0894*** |

**Table S1a.** The results of the ANOVA fits for the male CRF in each combination of female and background. Adjusted alpha rate for 4 comparisons is α = 0.01274

| *Response* | *Female* | *Background* | *Variable* | *df* | *F* | *P* |
|---|---|---|---|---|---|---|
| Fitness | CG | HC | Selection | ***1*** | ***29.4353*** | ***<0.0001*** |
| | | | Replicate | ***2*** | ***1.5767*** | ***0.22*** |
| | | | Sel:Rep | ***2*** | ***3.0820*** | ***0.0542*** |
| Fitness | Cr | HC | Selection | ***1*** | ***59.8226*** | ***<0.0001*** |
| | | | Replicate | ***2*** | ***3.1641*** | ***0.0567*** |
| | | | Sel:Rep | ***2*** | ***0.1099*** | ***0.90*** |
| Fitness | CG | WT | Selection | ***1*** | ***5.8642*** | ***0.0189*** |
| | | | Replicate | ***2*** | ***0.0822*** | ***0.92*** |
| | | | Sel:Rep | ***2*** | ***0.6021*** | ***0.55*** |
| Fitness | Cr | WT | Selection | ***1*** | ***0.5599*** | ***0.46*** |
| | | | Replicate | ***2*** | ***0.7995*** | ***0.46*** |
| | | | Sel:Rep | ***2*** | ***0.7617*** | ***0.48*** |

**Table S2.** The results of the ANOVA fit for the male CRF, for target animals with CG autosomes

| *Response* | *Variable* | *df* | *F* | *P* |
|---|---|---|---|---|
| Fitness | Selection | ***1*** | ***2.3793*** | ***0.13*** |
| | Replicate | ***2*** | ***0.5659*** | ***0.57*** |
| | Sel:Rep | ***2*** | ***0.4788*** | ***0.62*** |

**Table S3.** The results of the ANOVA fit for the male CRF, for target animals with a CG cytotype

| *Response* | *Variable* | *df* | *F* | *P* |
|---|---|---|---|---|
| Fitness | Selection | ***1*** | ***5.2034*** | ***0.03*** |
| | Replicate | ***2*** | ***0.0006*** | ***0.99*** |
| | Sel:Rep | ***2*** | ***2.0104*** | ***0.15*** |

**Table S4.** The results of the ANOVA fit for the male CRF, for target animals with a CG Y chromosome

| *Response* | *Variable* | *df* | *F* | *P* |
|---|---|---|---|---|
| Fitness | Selection | ***1*** | ***27.1966*** | ***<0.0001*** |
| | Replicate | ***2*** | ***12.5216*** | ***0.0002*** |
| | Sel:Rep | ***2*** | ***4.9401*** | ***0.015*** |

**Table S5.** The results of the ANOVA fit for the female CRF, for target animals with a CG cytotype

| *Response* | *Variable* | *df* | *F* | *P* |
|---|---|---|---|---|
| Fitness | Selection | ***1*** | ***3.9303*** | ***0.0517*** |
| | Replicate | ***2*** | ***13.8080*** | ***<0.0001*** |
| | Sel:Rep | ***2*** | ***3.5833*** | ***0.0334*** |

**Table S4a.** The results of the ANOVA fit for the male CRF, for target animals with a CG Y chromosome, separated by replicate. Adjusted alpha rate for 3 comparisons is α = 0.0169

| *Response* | *Replicate* | *Variable* | *df* | *F* | *P* |
|---|---|---|---|---|---|
| Fitness | 1 | Selection | ***1*** | ***24.840*** | ***0.0011*** |
| | 3 | Selection | ***1*** | ***27.085*** | ***0.0008*** |
| | 5 | Selection | ***1*** | ***0.1523*** | ***0.71*** |

**Table S5a.** The results of the ANOVA fit for the female CRF, for target animals with a CG cytotype, separated by replicate. Adjusted alpha rate for 3 comparisons is α = 0.0169

| *Response* | *Replicate* | *Variable* | *df* | *F* | *P* |
|---|---|---|---|---|---|
| Fitness | 1 | Selection | ***1*** | ***0.0074*** | ***0.93*** |
| | 3 | Selection | ***1*** | ***0.0363*** | ***0.85*** |
| | 5 | Selection | ***1*** | ***17.381*** | ***0.0004*** |

**Table S6.** The results of the fully factorial ANOVA fit for the male mating success.

| *Response* | *Variable* | *df* | *F* | *P* |
|---|---|---|---|---|
| Mating Success | Selection | ***1*** | ***11.852*** | ***0.0006*** |
| | Replicate | ***2*** | ***6.9319*** | ***0.0010*** |
| | Female | ***1*** | ***6.0212*** | ***0.0144*** |
| | Background | ***1*** | ***1.8860*** | ***0.17*** |
| | Sel:Rep | ***2*** | ***0.2892*** | ***0.75*** |
| | Sel:Fem | ***1*** | ***0.2098*** | ***0.65*** |
| | Sel:Bg | ***1*** | ***0.0339*** | ***0.85*** |
| | Fem:Bg | ***1*** | ***0.0978*** | ***0.75*** |
| | Fem:Rep | ***2*** | ***0.0025*** | ***0.99*** |
| | Bg:Rep | ***2*** | ***0.6487*** | ***0.52*** |
| | Sel:Fem:Bg | ***1*** | ***0.4053*** | ***0.52*** |
| | Sel:Bg:Rep | ***2*** | ***1.2380*** | ***0.29*** |
| | Sel:Fem:Rep | ***2*** | ***2.5206*** | ***0.081*** |
| | Fem:Bg:Rep | ***2*** | ***0.7242*** | ***0.48*** |
| | Sel:Fem:Bg:Rep | ***2*** | ***0.4756*** | ***0.62*** |

**Table S6a.** The results of the ANOVA fits for the male mating success in each replicate. Adjusted alpha rate for 3 comparisons is α = 0.0169

| *Response* | *Replicate* | *Variable* | *df* | *F* | *P* |
|---|---|---|---|---|---|
| Mating Success | 1 | Selection | ***1*** | ***5.7934*** | ***0.0166*** |
| | | Background | ***1*** | ***0.0895*** | ***0.76*** |
| | | Female | ***1*** | ***2.0137*** | ***0.16*** |
| | | Sel:Bg | ***1*** | ***1.3593*** | ***0.24*** |
| | | Sel:Fem | ***1*** | ***2.9241*** | ***0.0881*** |
| | | Bg:Fem | ***1*** | ***0.0427*** | ***0.84*** |
| | | Sel:Bg:Fem | ***1*** | ***0.4183*** | ***0.52*** |
| Mating Success | 3 | Selection | ***1*** | ***4.6478*** | ***0.0318*** |
| | | Background | ***1*** | ***2.9265*** | ***0.0880*** |
| | | Female | ***1*** | ***2.0929*** | ***0.15*** |
| | | Sel:Bg | ***1*** | ***0.2155*** | ***0.64*** |
| | | Sel:Fem | ***1*** | ***0.3033*** | ***0.58*** |
| | | Bg:Fem | ***1*** | ***1.3429*** | ***0.25*** |
| | | Sel:Bg:Fem | ***1*** | ***0.7699*** | ***0.38*** |
| Mating Success | 5 | Selection | ***1*** | ***1.9735*** | ***0.16*** |
| | | Background | ***1*** | ***0.1392*** | ***0.71*** |
| | | Female | ***1*** | ***1.9094*** | ***0.17*** |
| | | Sel:Bg | ***1*** | ***1.0175*** | ***0.31*** |
| | | Sel:Fem | ***1*** | ***2.0905*** | ***0.15*** |
| | | Bg:Fem | ***1*** | ***0.1635*** | ***0.69*** |
| | | Sel:Bg:Fem | ***1*** | ***0.1683*** | ***0.68*** |

**Table S7.** The results of the fully factorial ANOVA fit for the male mating latency.

| *Response* | *Variable* | *df* | *F* | *P* |
|---|---|---|---|---|
| log(latency+1) | Selection | ***1*** | ***4.8707*** | ***0.0277*** |
| | Replicate | ***2*** | ***2.4328*** | ***0.0887*** |
| | Female | ***1*** | ***0.0036*** | ***0.48*** |
| | Background | ***1*** | ***0.5040*** | ***0.95*** |
| | Sel:Rep | ***2*** | ***2.6575*** | ***0.0710*** |
| | Sel:Fem | ***1*** | ***0.3154*** | ***0.57*** |
| | Sel:Bg | ***1*** | ***0.1183*** | ***0.73*** |
| | Fem:Bg | ***1*** | ***2.4795*** | ***0.12*** |
| | Fem:Rep | ***2*** | ***1.7631*** | ***0.17*** |
| | Bg:Rep | ***2*** | ***0.2983*** | ***0.74*** |
| | Sel:Fem:Bg | ***1*** | ***0.5340*** | ***0.46*** |
| | Sel:Bg:Rep | ***2*** | ***0.1233*** | ***0.88*** |
| | Sel:Fem:Rep | ***2*** | ***2.0226*** | ***0.13*** |
| | Fem:Bg:Rep | ***2*** | ***0.2421*** | ***0.78*** |
| | Sel:Fem:Bg:Rep | ***2*** | ***2.0371*** | ***0.13*** |

**Table S8.** The results of the fully factorial ANOVA fit for the male mating duration.

| *Response* | *Variable* | *df* | *F* | *P* |
|---|---|---|---|---|
| Duration | Selection | ***1*** | ***5.7565*** | ***0.0168*** |
| | Replicate | ***2*** | ***3.9574*** | ***0.0197*** |
| | Female | ***1*** | ***13.122*** | ***0.0003*** |
| | Background | ***1*** | ***11.632*** | ***0.0007*** |
| | Sel:Rep | ***2*** | ***0.4611*** | ***0.63*** |
| | Sel:Fem | ***1*** | ***0.8901*** | ***0.34*** |
| | Sel:Bg | ***1*** | ***2.9102*** | ***0.0885*** |
| | Fem:Bg | ***1*** | ***0.0755*** | ***0.78*** |
| | Fem:Rep | ***2*** | ***0.0931*** | ***0.91*** |
| | Bg:Rep | ***2*** | ***0.3389*** | ***0.71*** |
| | Sel:Fem:Bg | ***1*** | ***1.0318*** | ***0.31*** |
| | Sel:Bg:Rep | ***2*** | ***0.1324*** | ***0.88*** |
| | Sel:Fem:Rep | ***2*** | ***0.0675*** | ***0.93*** |
| | Fem:Bg:Rep | ***2*** | ***0.2644*** | ***0.77*** |
| | Sel:Fem:Bg:Rep | ***2*** | ***1.4654*** | ***0.23*** |

**Table S8a.** The results of the ANOVA fits for the male mating duration in each replicate. Adjusted alpha rate for 3 comparisons is α = 0.0169

| *Response* | *Replicate* | *Variable* | *df* | *F* | *P* |
|---|---|---|---|---|---|
| Mating Duration | 1 | Selection | ***1*** | ***0.6031*** | ***0.44*** |
| | | Background | ***1*** | ***6.2255*** | ***0.0134*** |
| | | Female | ***1*** | ***7.2311*** | ***0.0077*** |
| | | Sel:Bg | ***1*** | ***2.4218*** | ***0.12*** |
| | | Sel:Fem | ***1*** | ***0.0859*** | ***0.77*** |
| | | Bg:Fem | ***1*** | ***0.5709*** | ***0.45*** |
| | | Sel:Bg:Fem | ***1*** | ***0.0740*** | ***0.78*** |
| Mating Duration | 3 | Selection | ***1*** | ***2.8546*** | ***0.0928*** |
| | | Background | ***1*** | ***1.4336*** | ***0.23*** |
| | | Female | ***1*** | ***3.3933*** | ***0.0671*** |
| | | Sel:Bg | ***1*** | ***0.3654*** | ***0.55*** |
| | | Sel:Fem | ***1*** | ***0.6404*** | ***0.42*** |
| | | Bg:Fem | ***1*** | ***0.1182*** | ***0.72*** |
| | | Sel:Bg:Fem | ***1*** | ***0.0285*** | ***0.86*** |
| Mating Duration | 5 | Selection | ***1*** | ***2.9956*** | ***0.085*** |
| | | Background | ***1*** | ***5.4032*** | ***0.0214*** |
| | | Female | ***1*** | ***3.0061*** | ***0.085*** |
| | | Sel:Bg | ***1*** | ***0.7247*** | ***0.39*** |
| | | Sel:Fem | ***1*** | ***0.1817*** | ***0.67*** |
| | | Bg:Fem | ***1*** | ***0.0225*** | ***0.88*** |
| | | Sel:Bg:Fem | ***1*** | ***3.5357*** | ***0.062*** |

**Table S9.** The results of the fully factorial ANOVA fit for the fecundity induced by males.

| *Response* | *Variable* | *df* | *F* | *P* |
|---|---|---|---|---|
| Fecundity induced | Selection | ***1*** | ***0.3654*** | ***0.55*** |
| | Replicate | ***2*** | ***4.3529*** | ***0.0133*** |
| | Female | ***1*** | ***782.20*** | ***<0.0001*** |
| | Background | ***1*** | ***148.61*** | ***<0.0001*** |
| | Sel:Rep | ***2*** | ***0.4006*** | ***0.67*** |
| | Sel:Fem | ***1*** | ***0.1327*** | ***0.72*** |
| | Sel:Bg | ***1*** | ***2.5480*** | ***0.11*** |
| | Fem:Bg | ***1*** | ***21.246*** | ***<0.0001*** |
| | Fem:Rep | ***2*** | ***1.2105*** | ***0.30*** |
| | Bg:Rep | ***2*** | ***0.3138*** | ***0.73*** |
| | Sel:Fem:Bg | ***1*** | ***0.0742*** | ***0.78*** |
| | Sel:Bg:Rep | ***2*** | ***1.4759*** | ***0.23*** |
| | Sel:Fem:Rep | ***2*** | ***0.0181*** | ***0.98*** |
| | Fem:Bg:Rep | ***2*** | ***0.1891*** | ***0.83*** |
| | Sel:Fem:Bg:Rep | ***2*** | ***1.7209*** | ***0.18*** |

**Table S9a.** The results of the ANOVA fits for the fecundity induced by target males in each combination of female and background. Adjusted alpha rate for 4 comparisons is α = 0.01274

| *Response* | *Female* | *Background* | *Variable* | *df* | *F* | *P* |
|---|---|---|---|---|---|---|
| Fecundity induced | CG | HC | Selection | ***1*** | ***3.3938*** | ***0.0678*** |
| | | | Replicate | ***2*** | ***3.1436*** | ***0.0465*** |
| | | | Sel:Rep | ***2*** | ***0.2288*** | ***0.79*** |
| Fecundity induced | Cr | HC | Selection | ***1*** | ***2.0226*** | ***0.16*** |
| | | | Replicate | ***2*** | ***2.3028*** | ***0.10*** |
| | | | Sel:Rep | ***2*** | ***2.6918*** | ***0.0719*** |
| Fecundity induced | CG | WT | Selection | ***1*** | ***0.0000*** | ***0.99*** |
| | | | Replicate | ***2*** | ***0.8594*** | ***0.42*** |
| | | | Sel:Rep | ***2*** | ***0.2003*** | ***0.82*** |
| Fecundity induced | Cr | WT | Selection | ***1*** | ***0.1394*** | ***0.71*** |
| | | | Replicate | ***2*** | ***0.8620*** | ***0.42*** |
| | | | Sel:Rep | ***2*** | ***0.4580*** | ***0.63*** |

**Table S10.** The results of the fully factorial ANOVA fit for the sex ratio of broods sired by target males.

| *Response* | *Variable* | *df* | *F* | *P* |
|---|---|---|---|---|
| Brood sex ratio | Selection | ***1*** | ***2.2747*** | ***0.15*** |
| | Replicate | ***2*** | ***0.1619*** | ***0.85*** |
| | Female | ***1*** | ***0.0126*** | ***0.91*** |
| | Background | ***1*** | ***0.3660*** | ***0.55*** |
| | Sel:Rep | ***2*** | ***0.4732*** | ***0.62*** |
| | Sel:Fem | ***1*** | ***1.1934*** | ***0.27*** |
| | Sel:Bg | ***1*** | ***0.3567*** | ***0.55*** |
| | Fem:Bg | ***1*** | ***0.0863*** | ***0.77*** |
| | Fem:Rep | ***2*** | ***0.8309*** | ***0.44*** |
| | Bg:Rep | ***2*** | ***1.3983*** | ***0.25*** |
| | Sel:Fem:Bg | ***1*** | ***0.1977*** | ***0.66*** |
| | Sel:Bg:Rep | ***2*** | ***0.7795*** | ***0.46*** |
| | Sel:Fem:Rep | ***2*** | ***1.0350*** | ***0.36*** |
| | Fem:Bg:Rep | ***2*** | ***0.0947*** | ***0.90*** |
| | Sel:Fem:Bg:Rep | ***2*** | ***0.7588*** | ***0.47*** |

**Table S11a.** The results of the fully factorial ANOVA fit on a generalized linear model (binomial error distribution) on the number of target males that sired 100% of their mate's offspring.

| *Response* | *Variable* | *df* | *F* | *P* |
|---|---|---|---|---|
| P2 part 1 | Selection | ***1*** | ***0.0158*** | ***0.90*** |
| | Replicate | ***2*** | ***0.3192*** | ***0.73*** |
| | Female | ***1*** | ***3.9988*** | ***0.0458*** |
| | Background | ***1*** | ***44.164*** | ***<0.0001*** |
| | Sel:Rep | ***2*** | ***0.9428*** | ***0.39*** |
| | Sel:Fem | ***1*** | ***1.3406*** | ***0.25*** |
| | Sel:Bg | ***1*** | ***0.0784*** | ***0.78*** |
| | Fem:Bg | ***1*** | ***27.517*** | ***<0.0001*** |
| | Fem:Rep | ***2*** | ***1.7691*** | ***0.17*** |
| | Bg:Rep | ***2*** | ***1.4932*** | ***0.22*** |
| | Sel:Fem:Bg | ***1*** | ***0.1202*** | ***0.73*** |
| | Sel:Bg:Rep | ***2*** | ***2.2303*** | ***0.11*** |
| | Sel:Fem:Rep | ***2*** | ***1.9455*** | ***0.14*** |
| | Fem:Bg:Rep | ***2*** | ***0.2614*** | ***0.77*** |
| | Sel:Fem:Bg:Rep | ***2*** | ***0.9112*** | ***0.40*** |

**Table S11b.** The results of the fully factorial ANOVA fit for offspring sired, by target males that did not sire 100% of their mate's offspring.

| *Response* | *Variable* | *df* | *F* | *P* |
|---|---|---|---|---|
| P2 | Selection | ***1*** | ***0.7596*** | ***0.38*** |
| | Replicate | ***2*** | ***0.2658*** | ***0.77*** |
| | Female | ***1*** | ***86.891*** | ***<0.0001*** |
| | Background | ***1*** | ***138.31*** | ***<0.0001*** |
| | Sel:Rep | ***2*** | ***0.2050*** | ***0.81*** |
| | Sel:Fem | ***1*** | ***0.3101*** | ***0.58*** |
| | Sel:Bg | ***1*** | ***0.4304*** | ***0.51*** |
| | Fem:Bg | ***1*** | ***87.723*** | ***<0.0001*** |
| | Fem:Rep | ***2*** | ***0.6061*** | ***0.55*** |
| | Bg:Rep | ***2*** | ***1.0976*** | ***0.34*** |
| | Sel:Fem:Bg | ***1*** | ***2.3676*** | ***0.12*** |
| | Sel:Bg:Rep | ***2*** | ***2.5523*** | ***0.08*** |
| | Sel:Fem:Rep | ***2*** | ***1.5697*** | ***0.21*** |
| | Fem:Bg:Rep | ***2*** | ***0.0424*** | ***0.96*** |
| | Sel:Fem:Bg:Rep | ***2*** | ***1.2971*** | ***0.28*** |

**Table S12.** The results of the ANOVA fit for mate harm by target males.

| *Response* | *Variable* | *df* | *F* | *P* |
|---|---|---|---|---|
| Mortality | Selection | ***1*** | ***17.648*** | ***<0.0001*** |
| | Replicate | ***2*** | ***1.7037*** | ***0.19*** |
| | Background | ***1*** | ***18.967*** | ***<0.0001*** |
| | Sel:Rep | ***2*** | ***1.0799*** | ***0.34*** |
| | Sel:Bg | ***1*** | ***0.2510*** | ***0.62*** |
| | Bg:Rep | ***2*** | ***1.8731*** | ***0.16*** |
| | Sel:Bg:Rep | ***2*** | ***1.0710*** | ***0.35*** |
| Productivity | Selection | ***1*** | ***21.924*** | ***<0.0001*** |
| | Replicate | ***2*** | ***0.6079*** | ***0.55*** |
| | Background | ***1*** | ***164.21*** | ***<0.0001*** |
| | Sel:Rep | ***2*** | ***0.0897*** | ***0.91*** |
| | Sel:Bg | ***1*** | ***32.504*** | ***<0.0001*** |
| | Bg:Rep | ***2*** | ***0.5131*** | ***0.60*** |
| | Sel:Bg:Rep | ***2*** | ***1.6544*** | ***0.20*** |

**Figure S1.** (a) ML selection breeding design. (b) Recombination box design. CG females are denoted as DTW or DTP based on presence or absence of *bw*$^{D}$ marker.

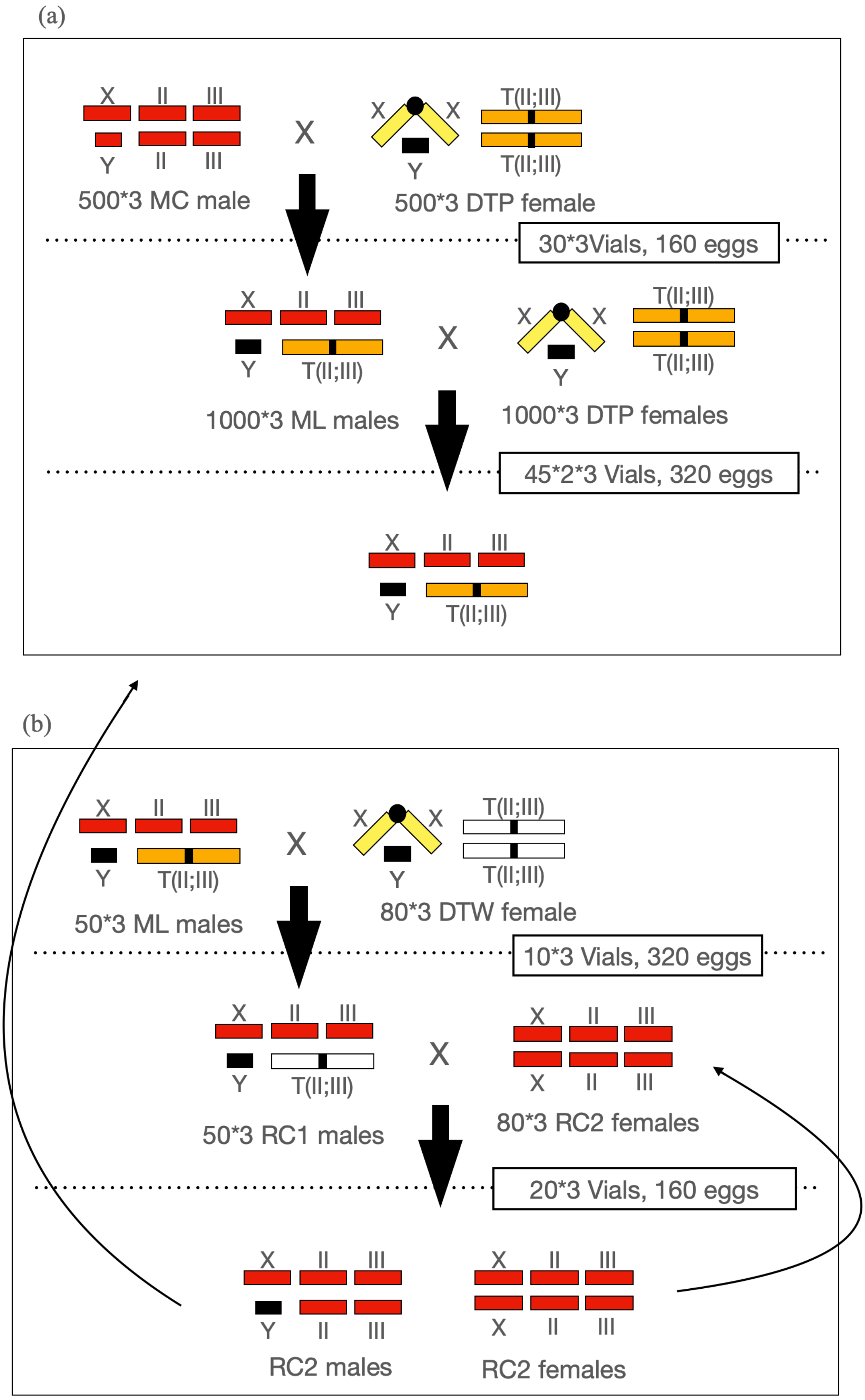

**Figure S2.** (a) HC males and WT males used for (b) CRF assays and (c) mating success and sperm offense assays. Females denoted as DTP in the schematic represent clone-generator (CG) females carrying translocated autosomes marked with recessive eye-colour markers. Where indicated, females denoted as DTW instead represent CG females carrying translocated autosomes marked with dominant eye-colour markers.

(a)

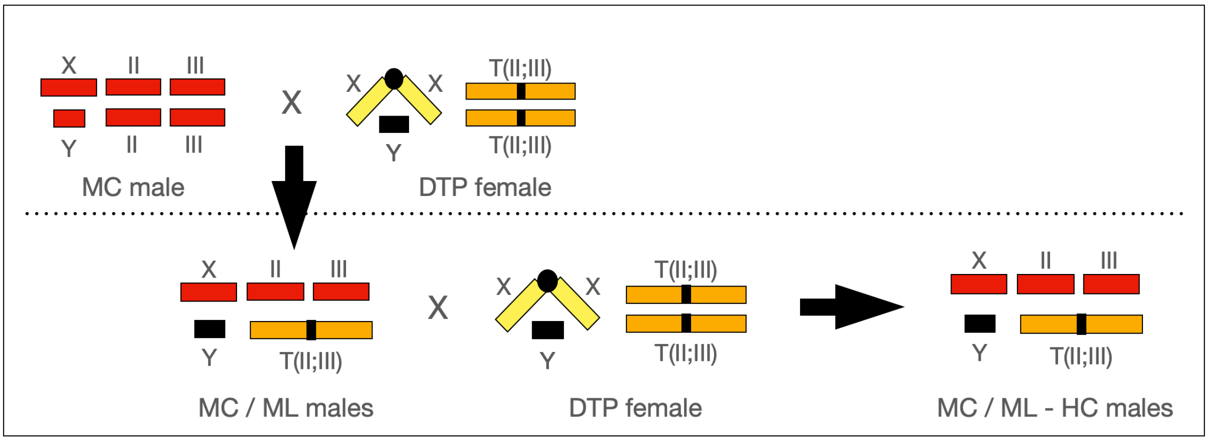


(b)

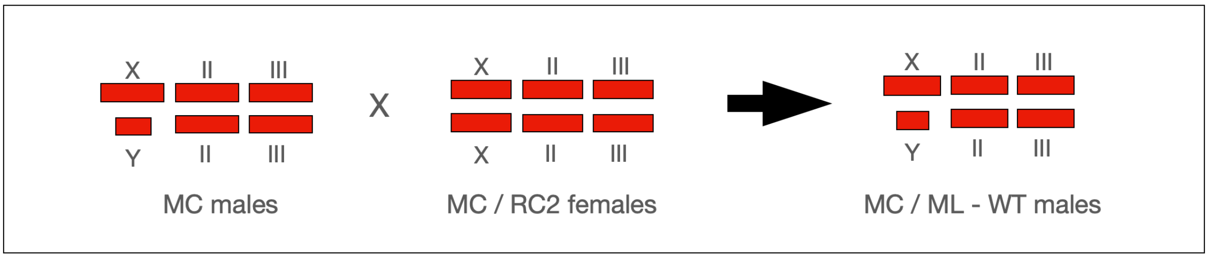

**Figure S3.** Single artefact males: (a) CG cyto, (b) CG Y, and (c) CG auto animals.

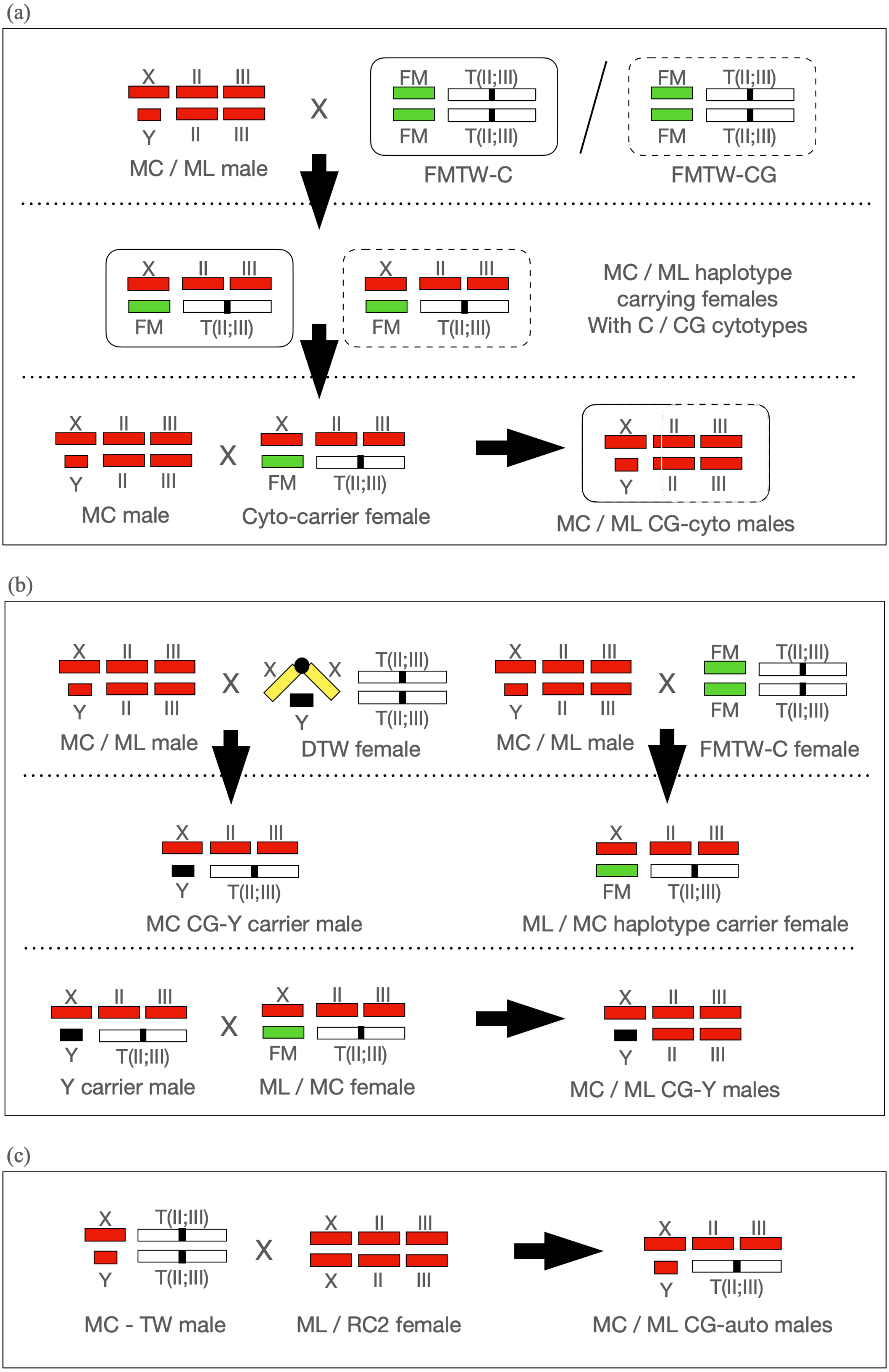

**Figure S4.** Single artefact females – (a) CG cyto and (b) CG auto females

(a)

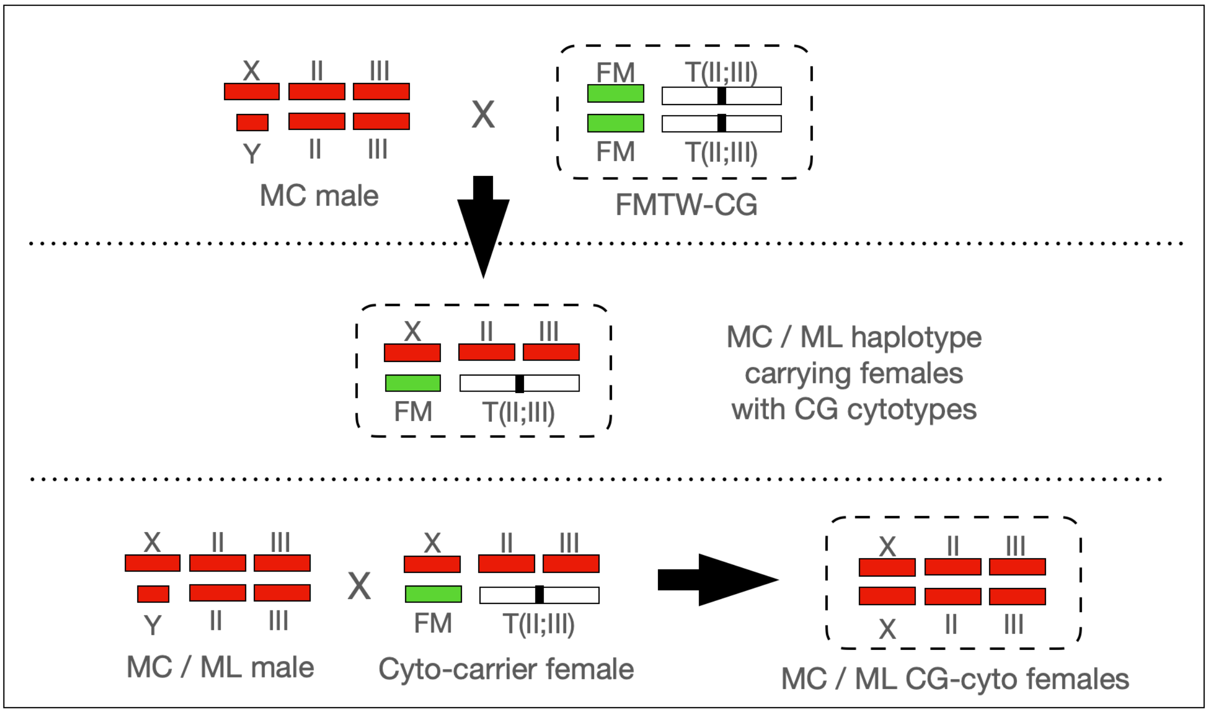


(b)

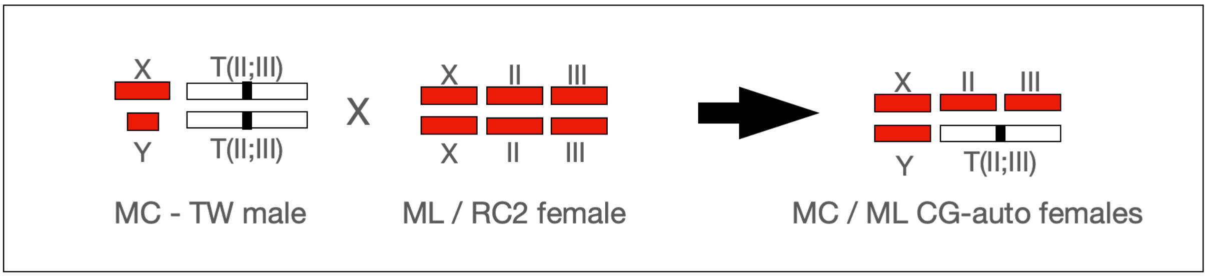
